# Synthetic genome rearrangement reveals dynamics of chromosome evolution shaped by hierarchical chromatin organization

**DOI:** 10.1101/2021.07.19.453002

**Authors:** Sijie Zhou, Yi Wu, Yu Zhao, Zhen Zhang, Limin Jiang, Lin Liu, Yan Zhang, Jijun Tang, Ying-Jin Yuan

## Abstract

Synthetic genome evolution provides a dynamic approach to systematically and straightforwardly explore evolutionary processes. SCRaMbLE is an evolutionary system intrinsic to the synthetic yeast genome that can rapidly drive structural variations. Here, we detect over 260,000 rearrangement events after SCRaMbLEing of a novel yeast strain harboring 6 synthetic yeast chromosomes. Remarkably, we find that the rearrangement events exhibit a specific landscape of rearrangement frequency. We further reveal that the landscape is shaped by combinatorial effects of chromatin accessibility and spatial contact probability. The rearrangements tend to occur in 3D spatially proximal and chromatin-accessible regions. Enormous numbers of rearrangements by SCRaMbLE provide a driving force to potentiate directed genome evolution, and investigation of the rearrangement landscape offers mechanistic insights into the dynamics of genome evolution.

## Introduction

Studying the processes and mechanisms of genome evolution is critical to understanding genetic diversity and species diversity at the genomic level^1-4^. *S. cerevisiae*, a powerful model organism for eukaryotic genome evolution^5-7^ has been subjected to comparative genomic studies that provided mechanistic insights^8-12^. These studies relied on relatively static genomic sequences and may have missed many details of dynamic processes. Synthetic genomes, assembled from scratch and incorporated with a variety of designer features, have greatly facilitated the study of genome evolution and engineering, such as genome minimization^13^, genetic codon recoding^14, 15^, introduction of synthetic parts^16^ and data storage^17, 18^. SCRaMbLE with symmetrical loxP sites (loxPsym) positioned in the 3′ untranslated regions (3′ UTRs) of all nonessential genes in the Synthetic Yeast Genome Project (Sc2.0) has also been used in the study of genome evolution recently^16, 19^. Induced Cre recombinase activity quickly triggers recombination between loxPsym sites and generates structural variations, including deletions, inversions, duplications, and translocations^20-27^.

In this study, we first consolidated six individual synthetic chromosomes (synII, synIII, synV, synVI, synIXR and synX) into a single haploid strain with an orthogonal site-specific recombination system, which enabled the eliminations of counterpart native chromosomes^19, 28-32^. In this strain with nearly half a synthetic genome, we induced SCRaMbLE and generated massive rearrangement events with both intra- and inter-chromosomal recombination. We comprehensively sequenced this SCRaMbLEd pool and detected over 260,000 rearrangement events via a loxPsym junction analysis method. By analyzing each rearrangement event, we uncovered a stable rearrangement landscape for synthetic chromosomes that was correlated to local chromatin structures and three-dimensional genome architecture analyzed via Assay for Transposase-Accessible Chromatin sequencing (ATAC-seq) and genome-wide chromosome conformation capture (Hi-C). Our study provides insight into the combinatorial effect of hierarchical chromatin organization on the dynamics of genome evolution.

## Results

### Consolidation and SCRaMbLE of six synthetic chromosomes

To build the strain with multiple synthetic chromosomes, we used a stepwise method, starting with six individual yeast strains that contain single synthetic chromosomes (synII, synIII, synV, synVI, synIXR, synX). First, we obtained two haploid strains of opposite mating types, yYW169 (synV, synX) and yZY192 (synII, synIII, synVI and synIXR), using endoreduplication intercrossing^16^. To accelerate the consolidation, we developed a new strategy using chromosome elimination by Vika/vox, a site-specific recombination system orthogonal to Cre/loxP (Fig. 1a)^33, 34^. First, we built a heterozygous diploid strain yYW268 by mating yYW169 and yZY192. The native chromosomes (II, III, V, VI, IX and X) were eliminated successively, with their centromeres excised by vox recombination (Supplementary Fig.1). Following endoreduplication, sporulation and tetrad dissection, one haploid strain (yYW394) with the six aforementioned native chromosomes swapped with synthetic chromosomes was obtained, comprising ∼2.61 Mb (∼22.0% of the yeast genome) (Supplementary Table 1). Its karyotype and genome sequence were confirmed by pulsed-field gel electrophoresis and Whole Genome Sequencing (WGS) (Supplementary Fig.2b, d). The yYW394 strain grew robustly at 30 □ but exhibited a growth defect at 37 □ (Supplementary Fig.2c). To recover its fitness to wild-type levels, adaptive laboratory evolution of yYW394 was performed and generated a new healthy strain yZSJ025, which was used for further analysis (Supplementary Fig.2).

**Fig.1.**
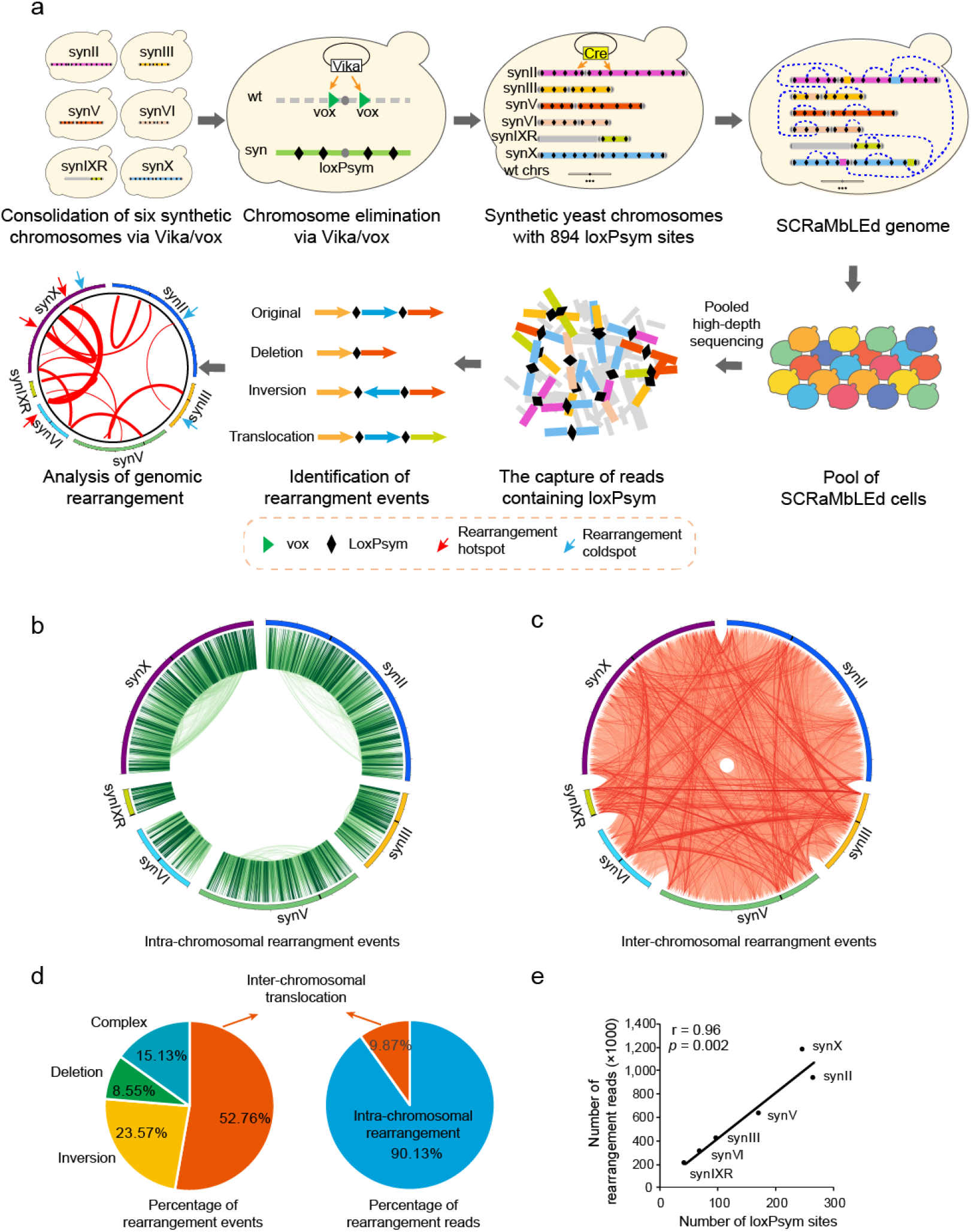
Consolidation and SCRaMbLE of six synthetic chromosomes. **a**, Schematic diagram showing the construction of the synthetic yeast strain yZSJ025 and the analysis of rearrangement events following SCRaMbLE. **b**, Circos plot of intra-chromosomal rearrangement events in yZSJ025. **c**, Circos plot of inter-chromosomal rearrangement events in yZSJ025. In (a) and (c), two loxPsym sites where rearrangement events occur are connected with a line, with its intensity representing the frequency of this event. **d**, Classifications of rearrangement events with percentages of different groups calculated by event numbers and read numbers respectively. Different groups of events are labeled in different colors as indicated. **e**, Correlation analyses between the rearrangement frequency and the number of loxPsym sites on each chromosome. Pearson correlation analysis was applied to determine the correlation coefficient and associated p values.

A Cre recombinase expression plasmid pYW085 (pRS413-pCLB2-Cre-EBD) was transformed into yZSJ025, comprising 894 loxPsym sites (Supplementary Fig.3). SCRaMbLE was induced by addition of β-estradiol. The SCRaMbLEd cells were then diluted in fresh YPD liquid medium without β-estradiol (Fig. 1a), and subjected to deep sequencing (∼600,000×). All reads (150 bp each) were screened for the presence of loxPsym sequence, then aligned to the reference sequence of original synthetic chromosomes. Those with flanking sequences different from the references, hereafter called rearrangement reads, were then used to identify and classify rearrangement events^25^ (Supplementary Fig.4). Identical reads were considered as one rearrangement event. Totally, 263,520 rearrangement events, including 124,499 (47.24%) intra- and 139,021 (52.76%) inter-chromosomal events were detected (Fig. 1b, c). We further analyzed the intra-chromosomal rearrangement events and found 62,106 (23.57%) inversions, 22,526 (8.55%) deletions and 39,867 (15.13%) complex rearrangement events, such as duplication and circularization (Fig. 1d and Supplementary Fig.4). We also counted the reads from each rearrangement. The numbers of identical reads represented the frequencies of corresponding rearrangement events. The inter-chromosomal recombination only accounted for 9.87% of total read numbers, indicating that these are relatively low frequency events compared to intra-chromosomal rearrangements. We next investigated whether there is any chromosome preference for rearrangements. To answer this question, we plotted the number of rearrangements for each chromosome against the number of loxPsym sites in that chromosome. As expected, the number of rearrangements correlates with the number of loxpsym sites per chromosome (r = 0.96, *p* = 0.002) (Fig. 1e).

### A specific rearrangement pattern of the synthetic yeast chromosomes

In theory, SCRaMbLE can generate rearrangements between any two loxPsym sites on synthetic chromosomes. However, different chromosomal loci showed high or low rearrangement frequencies (Fig. 1b, c). To estimate the local rearrangement frequencies along the synthetic chromosomes in our SCRaMbLEd pool, we counted the number of rearrangement reads from each loxPsym site, generating a landscape of rearrangement frequencies (Fig. 2a). Notably, rearrangement reads were identified from all 877 loxPsym sites, indicating the sequencing coverage was sufficient to cover all the loci. The loxPsym sites differed from each other in rearrangement frequencies, ranging from 1,027 to 44,353 per site. We selected 90 loxPsym sites (∼10%) with the highest or lowest rearrangement frequencies as hotspots or coldspots, respectively, for further analysis. Rearrangement frequencies of hotspots were higher than 17,800 per site while for coldspots, the frequencies were lower than 3,400 per site. These rearrangement frequencies were significantly different from the average frequency of all loxPsym sites (Fig. 2b). Interestingly, the rearrangement frequency landscapes of independent SCRaMbLE experiments were highly reproducible for all the synthetic chromosomes, indicating certain areas of the genome have the potential to undergo rearrangement at higher frequency (Supplementary Figs.5, 6).

**Fig.2.**
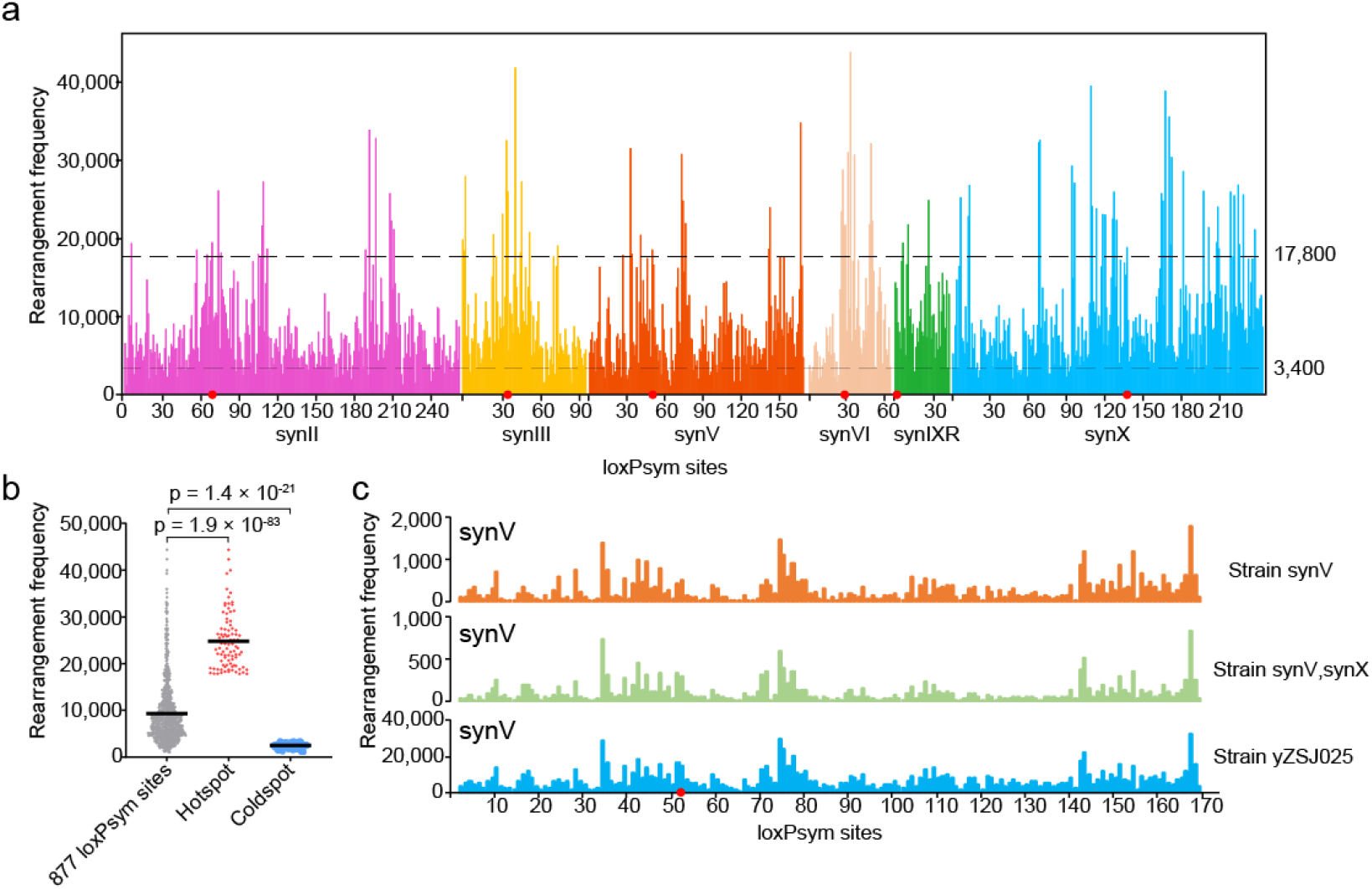
Investigating rearrangement frequencies revealed a specific rearrangement pattern among synthetic yeast chromosomes. **a**, Landscape of rearrangement frequencies along synII, synIII, synV, synVI, synIXR and synX. 17 loxPsym sites were initially inserted at the synthetic telomere regions, with flanking sequences not distinguishable from each other. They were excluded from the identification of rearrangements reads. **b**, Comparison of rearrangement frequencies between hotspots and coldspots. Rearrangement frequencies of hotspots, coldspots and all 877 loxPsym sites. The points represent the rearrangement frequencies of each loxPsym site. Horizontal lines indicate weighted means. The *p* values were calculated using two-tailed paired two-sample *t*-tests. **c**, Intra-chromosomal rearrangement patterns of synV following SCRaMbLE in three different yeast strains, yXZX846 (synV), yYW169 (synV, synX) and yZSJ025 respectively. Pearson correlation analysis was applied to determine the correlation coefficient and associated p values. For (a) and (c), red dots indicate centromeres.

We also compared rearrangement patterns of the same synthetic chromosome in separate strains containing different numbers of synthetic chromosomes. Similar intra-chromosomal rearrangement patterns were observed for synV in the three synV-containing strains (yXZX846, yYW169 and yZSJ025) (Fig. 2c and Supplementary Fig 7a-c). The same result was also observed for synX in the two synX-containing strains (yYW169 and yZSJ025) (Supplementary Fig.7d, e). Overall, our results demonstrate that the patterns of SCRaMbLE followed the same biological reason on each synthetic chromosome with reproducible rearrangement hotspots and coldspots.

### Rearrangement frequency correlates with chromatin accessibility

To determine if the specific rearrangement patterns we observe are due to an underlying biological mechanism, we further investigated the genomic landscape. SCRaMbLE requires physical interaction between Cre recombinase and loxPsym sites, suggesting the importance of chromatin accessibility in determining rearrangement frequency in SCRaMbLEd cells. To test this hypothesis, we measured genome-wide chromatin accessibility in yZSJ025 strain by ATAC-seq^35^. The ATAC-seq signals from the window containing 400 bp upstream and downstream of each loxPsym site were collected and processed. The signals from each rearrangement hotspot and coldspot were normalized with the average signals of all 877 loxPsym sites. Interestingly, statistical analysis showed that the average ATAC-seq signals of hotspots were significantly higher, while coldspot signals were significantly weaker (Fig. 3a). The rearrangement frequencies and ATAC-seq signals of a typical loxPsym hotspot and coldspot from synX are shown in Fig. 3b. As a coldspot, the loxPsym site in *SET4* 3’ UTR had a weak ATAC-seq signal, while as a hotspot, the loxPsym site in *PRY3* 3′ UTR showed a strong ATAC-seq signal. In addition, we also observed that nucleosome occupancy was relatively low in the hotspots and high in the coldspots (Supplementary Fig.8). Both ATAC-seq and nucleosome occupancy indicated that chromatin accessibility is critical for determining patterns of rearrangement frequencies.

**Fig. 3.**
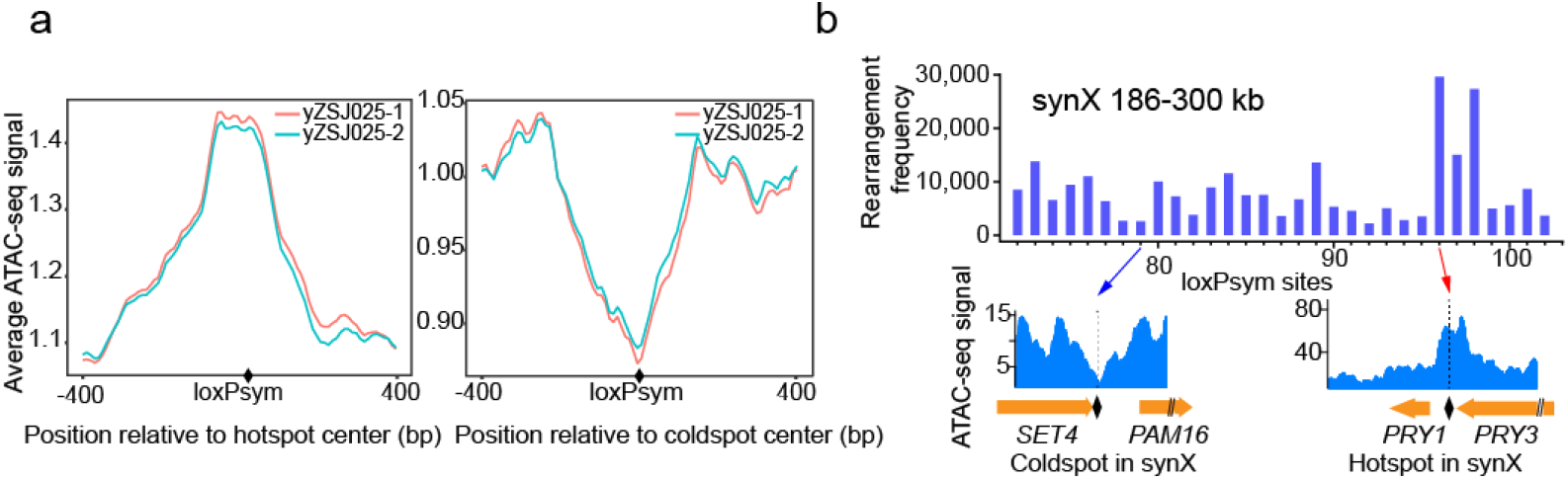
Rearrangement frequency in correlation with chromatin accessibility. **a**, Average ATAC-seq signals of the hotspot- and coldspot-centered 800 bp regions. **b**, ATAC-seq signals for a typical rearrangement hotspot and a coldspot in the 186-300 kb region of synX.

We also aligned the hotspots and coldspots to the corresponding region in native chromosomes. Using the same approach as before, we extracted and analyzed pre-published ATAC-seq data from wild-type yeast ^35^ (Supplementary Fig.9). ATAC-seq signals peaked at the positions of the hotspots, and remained weak at the coldspots in all six chromosomes, suggesting that sequence modifications of Sc2.0 do not perturb chromatin accessibility.

### Rearrangement frequency also correlates with 3D chromosome conformation

Next, we aimed to explore whether rearrangement frequency is correlated with the spatial proximity of loxPsym sites. Our SCRaMbLE system provided a unique platform to statistically evaluate the role of spatial proximity in chromosomal rearrangements. A genomic chromosome conformation capture approach (Hi-C) was thus carried out for yZSJ025, generating a contact map with the frequencies of spatial contacts between any two genomic loci (Supplementary Fig.10). We extracted contact frequencies from the synthetic chromosomes (Fig. 4a), and plotted rearrangement frequencies in a similar heatmap to facilitate direct comparison (Fig. 4b). The regions near the diagonal represent proximal loci in each chromosome. Consistent with this, these intra-chromosomal regions were the “hottest” in the Hi-C map (Fig. 4a), and they also exhibited the highest rearrangement frequency (Fig. 4b). Rearrangements tended to occur most frequently between adjacent loxPsym sites for all synthetic chromosomes as shown by statistical analysis of all rearrangement reads (Supplementary Fig.11).

**Fig.4.**
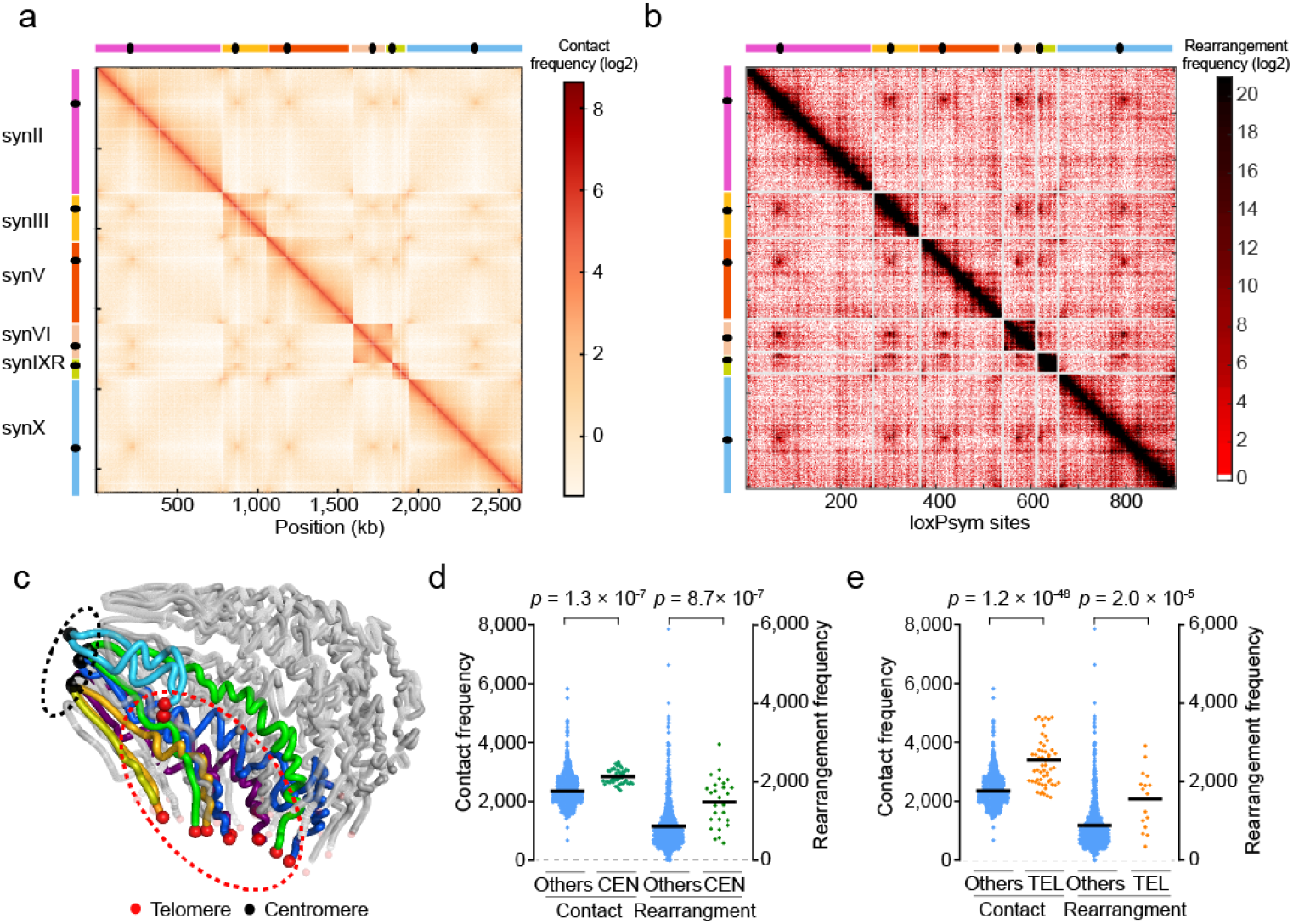
Rearrangement frequency in correlation with 3D chromosome conformation. **a**, Hi-C heatmap. The heatmap value for a site (*i, j*) is the contact probability (log2) between genomic loci *i* (horizontal axis) and *j* (vertical axis). Both axes are displayed at a 1 kb resolution. Spots with different heatmap values from low to high are colored from light yellow to red as indicated. **b**, Rearrangement frequency heatmap. The heatmap value for a site (*i, j*) is the rearrangement frequency (log2) of the event occurring at the loci *i* (vertical axis) and *j* (horizontal axis). Spots with different heatmap values are labeled in red with different intensities as indicated. **c**, 3D structures of synthetic chromosomes inferred from the Hi-C contact map displayed in panel (a). Synthetic chromosomes are labeled with different colors and the native chromosomes are shown in grey. Centromeres and telomeres were represented by dark and red spheres respectively. **d**, Comparisons of the contact probability and frequency of rearrangement events at loci in pericentromeric regions (CEN) and in other regions (Others). The CEN are centromere-centered 10 kb regions. **e**, Comparisons of the contact probability and frequency of rearrangement events at loci in peritelomeric regions (TEL) and in other regions (Others). The TEL are ≤5 kb from telomeres. For (d) and (e), each data point represents 1 kb bin in regions as indicated. Horizontal lines indicate weighted means. The *p* values were calculated using two-tailed paired two-sample *t*-tests.

Notably, the inter-chromosomal contact probability in the regions of centromeres and telomeres was obviously higher than that in other regions (Fig. 4a), consistent with centromeres being clustered around the spindle pole body and telomeres being clustered with the nuclear envelope for both wild-type^36^ and synthetic chromosomes^37^ (Fig. 4c). Compared to other regions, pericentromeric regions exhibited higher inter-chromosomal rearrangement frequency (Fig. 4b, d). Similar results were found for the peritelomeric regions of the synthetic chromosomes (Fig. 4b, e). Taken together, our results suggested that rearrangement events are generally more likely to occur between genomic loci in spatial proximity.

### Combinatorial effects of chromatin accessibility and chromosome conformation

As both chromatin accessibility and spatial proximity influence rearrangement frequency, we hypothesized that they have combinatorial effects on yeast chromosome organization. We tested our proposed model based on individual rearrangement events. If two loxPsym sites have similar chromatin accessibility (Fig. 5a), the frequency of rearrangement events was determined by their spatial proximity (Fig. 5b). For loxPsym sites with similar spatial proximity (Fig. 5c), the rearrangement frequency was mainly affected by their chromatin accessibility (Fig. 5d). Most interestingly, if two loxPsym sites had open chromatin but were distant in space, they might have a similar recombination frequency compared to other pairs of loxPsym sites that were proximal but buried in closed chromatin structures (Fig. 5e, f). In other words, the effect of chromatin accessibility on rearrangement frequency could be compensated for by the difference of spatial proximity, and *vice versa*. More generally, chromatin architecture and spatial conformation in the yeast genome affected the rearrangement frequency in a combinatorial manner during SCRaMbLE.

**Fig.5.**
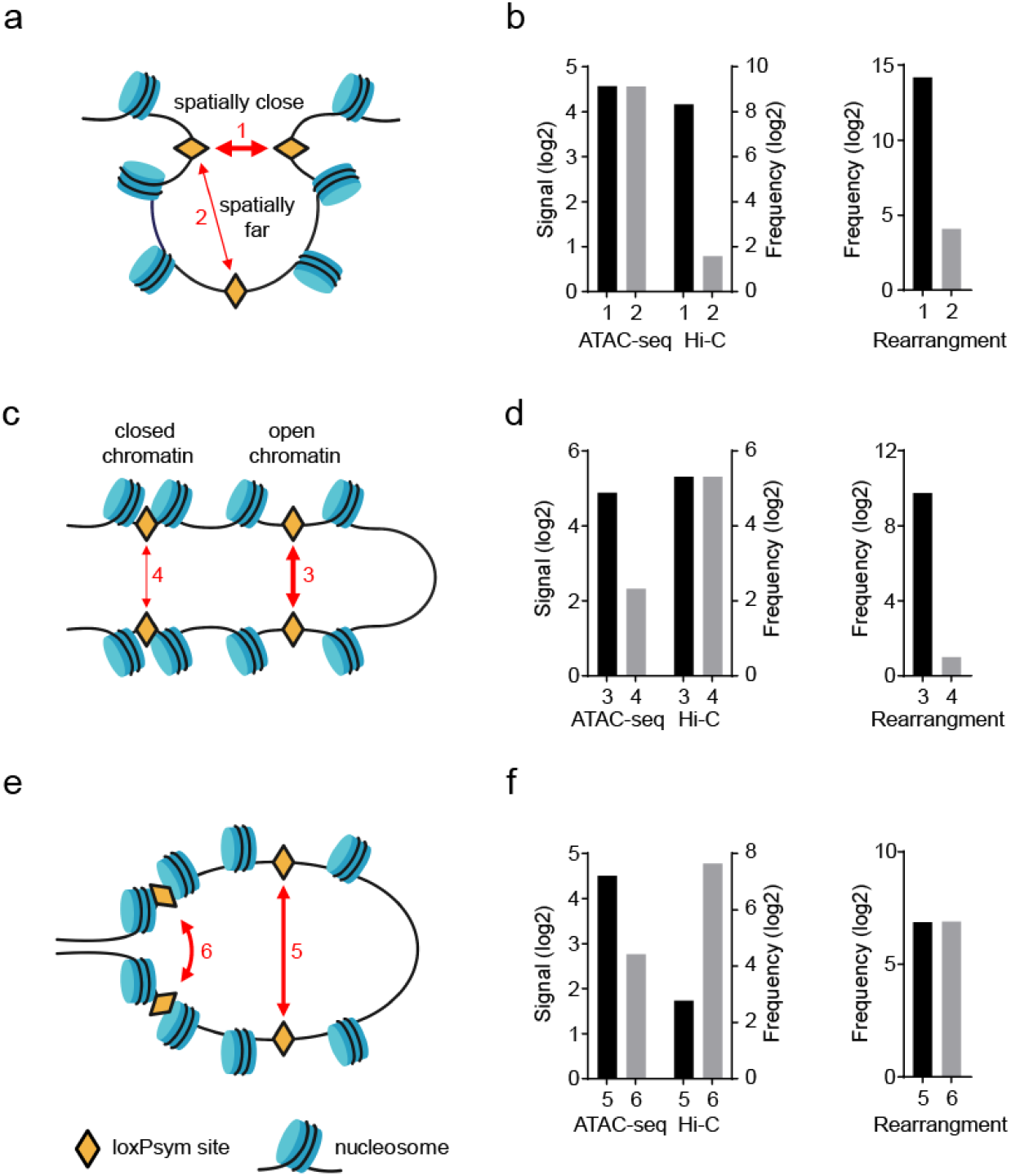
Mechanistic models of the effects of hierarchical chromatin organization on chromosomal rearrangement. **a**, A schematic model interpreting the difference of the frequencies of two rearrangement events with different contact probability. **b**, The loxPsym positions of pair 1: synX 183 kb and synX 184 kb; pair 2: synX 184 kb and synX 342 kb. **c**, A schematic model interpreting the difference of the frequencies of two rearrangement events with different chromatin accessibility. **d**, The loxPsym positions of pair 3: synII 186 kb and synII 204 kb; pair 4: synV 346 kb and synV 384 kb. **e**, A schematic model interpreting two rearrangement events with similar frequencies as a result of counter-effects by contact probability and chromatin accessibility. **f**, The loxPsym positions of pair 5: synVI 125 kb and synVI 202 kb; pair 6: synII 504 kb and synII 507 kb.

## Discussion

In this study, we induced SCRaMbLE in a novel strain with multiple synthetic chromosomes. Over 260,000 rearrangement events were detected in the SCRaMbLEd pool, covering all 894 loxPsym sites. This finding revealed the tremendous plasticity of the yeast genome. Previous studies of SCRaMbLE were mainly focused on the correlation of chromosomal rearrangement with nucleotide sequence^21, 23, 25, 27, 38^. Here, we discovered that the rearrangement frequency landscape was molded by chromatin accessibility and spatial proximity from an epigenetic perspective.

The biochemical essence of SCRaMbLE is recombination between pairs of different loxPsym sites in synthetic chromosomes catalyzed by Cre recombinase^39^. We speculated that local chromatin structure affected rearrangement by acting on the accessibility of Cre to loxPsym sites and that 3D chromosome conformation affected the contact probability of any two loxPsym sites. The use of synthetic chromosomes and SCRaMbLE allowed us to statistically reveal that inter-chromosomal rearrangement hotspots strikingly clustered at the peritelomeric and pericentromeric regions. Our results are consistent with previous reports that increased inter-chromosomal reshuffling occurred in the peritelomeric regions of domesticated and wild yeast isolates during genome evolution^8, 11^. The peritelomeric regions are functionally enriched for genes involved in secondary metabolism and stress responses that contribute to environmental adaptation^40-43^. High frequency of rearrangement in these regions could thus be an important driving force for evolution. Besides, inter-chromosomal rearrangements would result in swapping of chromosomal arms between two chromosomes. Recent genomic studies demonstrated that such rearrangements can cause reproductive isolation and promote incipient speciation^44-46^. Taking the combinatorial effect of chromatin accessibility and chromosome conformation into consideration, we speculated that chromatin structures might also play an important role in genome evolution in terms of the effects on the rearrangements in peritelomeric and pericentromeric regions.

In eukaryotic cells, altered 3D genome organization together with chromosomal structural variations have also been shown to be oncogenic, as in breast or ovarian cancers^47-49^. Most recently, a synthetic genomics approach was utilized to study spatial regulatory elements during embryonic development by reconstituting the *HoxA* cluster and many variants^50^. Our SCRaMbLE system, with multiple synthetic chromosomes, may also provide a unique platform to enable experimental characterization and a more mechanistic study of the role of genome rearrangement in disease.

All 16 synthetic yeast chromosomes and the novel tRNA neochromosome will be consolidated by chromosome swapping to build the final Sc2.0 strain^16, 28, 51^. The flexible and controllable synthetic yeast genome provides a unique model to systematically interrogate and explore the dynamics of eukaryotic genome evolution. The investigation of a large number of rearrangements indicated the tremendous plasticity of the yeast genome and the importance of hierarchical chromatin organization to the regional rates of chromosomal variation during genome evolution. These findings provide crucial insights that the variability and complexity of synthetic genome design can be further increased. Meanwhile, the influence of chromatin organization needs to be considered in the design and engineering of higher organism genomes.

## Materials and Methods

### Strains and plasmids

The plasmids and yeast strains used in this study were listed in Supplementary Table 4.

### Synthetic chromosome sequences and versions

The haploid draft strain yYW394 contains multiple synthetic chromosomes of synII (chr02_9_03), synIII (chr03_9_02), synV (chr05_9_04), synVI (chr06_9_03), synIXR (genebank JN020955), and synX (chr10_9_01) as published before^19, 28-32^. The final strain yZSJ025 contains the same synthetic chromosomes except some variants to improve its fitness, as listed in Supplementary Table 2. The genome sequencing data was submitted to NCBI Sequence Read Archive (SRA) under accession number PRJNA705059.

### Plasmid circuits construction

All plasmids were constructed using standard molecular cloning techniques and transformed into *E. coli* strain TOP 10 with standard protocols. Plasmid constructs were verified by restriction digests and Sanger sequencing by Genewiz. Restriction endonucleases and Phusion PCR kits were used from New England BioLabs. The KanMX-vox-CEN3-vox and hphNT1-vox-CEN3-vox were synthesized from Genewiz. (The distance between left vox and CENs is 100bp; the distance between right vox and CENs is 123bp). The 1kb homologous segments near the centromere of wild-type chromosome were PCR amplified from genome of BY4741. The wtII-hphNT1-vox-CEN3-vox, wtV-hphNT1-vox-CEN3-vox, wtVI-hphNT1-vox-CEN3-vox, wtIII-KanMX-vox-CEN3-vox, wtIX-KanMX-vox-CEN3-vox, and wtX-KanMX-vox-CEN3-vox were assembled with Gibson assembly into pUC19 linearized by *Sal*I and *Bam*HI digestion^52^.

### Yeast transformation

Yeast transformations in this study were performed with LiAc/SS/PEG method^53^. Briefly, the overnight YPD saturate culture of yeast cells were diluted into 5 mL fresh YPD medium for another 4-6 h culture starting from A_600_ of 0.2 at 30 °C in a shaker incubator at 220 rpm. Cells were washed with ddH_2_O followed by 0.1 M LiAc, and chilled on ice. 620 μL of 50 % polyethylene glycol (PEG-3350), 40 μL of salmon sperm DNA (100 mg/mL), 90 μL of 1 M LiAc, and 150 μL plasmid solution (100 ng DNA) were added in order to the cell pellet. The mixture was vortexed briefly to resuspend the cells and incubated at 30 °C for 30 min. 90 μL of dimethyl sulfoxide (DMSO) was then added to the mixture and a heat-shock for 18 min at 42 °C was performed. Cells were then collected and resuspended with 5 mM CaCl_2_. 100 μL of the cell suspension was then plated on SC–His for the selection of transformants. YPD, yeast extract peptone dextrose; SC, synthetic complete medium.

### Yeast mating process

Yeast strains of opposite mating types were incubated in selective medium overnight. 500 μL of each culture was washed with ddH_2_O, and the pellets were resuspended with 3 mL of YPD, and incubated at 30 °C in a shaker incubator at 220 rpm for 8 h. 100 µL of the mating suspension was subsequently sprayed on a selective medium agar plate for single colonies after incubation at 30 °C for 2 days. PCR screening for the MAT locus were performed to select diploid colonies.

### PCRTag assay for native chromosome elimination

PCRTags were landmarks in synthetic chromosomes including the peritelomeric and pericentromeric regions^16^. The loss of wild-type PCRTags associated with native chromosomes and the retained synthetic PCRTags on synthetic chromosomes were validated by colony PCR to assess the elimination of wtII, III, V, VI, IX, and X, using the primers listed in Supplementary Table 5.

### Construction of a poly-synthetic strain using chromosome elimination with Vika/vox

WtII-hphNT1-vox-CEN3-vox and wtIX-KanMX-vox-CEN3-vox constructs were linearized by *Sal*I and *Bam*HI, and integrated into the centromere regions of native chromosomes II and IX in the strain with synV and synX, which were then mated with the strain with synII, synIII, synVI and synIXR. Heterozygous diploid strains were transformed with a pGAL-Vika plasmid with *LEU2* auxotrophic marker^33^, and then inoculated in liquid YP+galactose (2%) medium for 12-18 h at 30 °C to turn on the Vika/vox recombination system. The entire native chromosome II and IX were eliminated after Vika recombinase expression was induced. 200 μL of culture was spread on plates with SC–Leu with dextrose medium. After incubation for 2 days at 30 °C, the single colonies were replicated to the plates with YPD+G418 (200ug/mL) and SC+Hygromycin B (200ug/mL) medium, to identify the colonies lost with native chromosome II and IX. Using PCRtag assays described above. Following the same strategy, wtV, wtX, wtVI, and wtIII were eliminated sequentially, generating a 2n–6 strain.

### Meiosis and sporulation

The 2n–6 strain was first transformed with a plasmid containing *MAT****a*** and *HIS3* marker. Yeasts were incubated in SC–His medium overnight at 30 °C with shaking at 220 rpm, after which 200 µL of the culture was transferred to 3 mL of YPD and then incubated at 30 °C with shaking at 220 rpm for 14 h for early stationary phase. The cells were washed with sterile ddH_2_O three times. 1 µL of 50× sporulation medium, 500 µL of the required amino acids (uracil, leucine), and 150 µL of 10% yeast extract were mixed together and then diluted with sterile ddH_2_O to a volume of 50 mL. All washed cells were transferred into sporulation solution and mixed well, which were subjected to sporulation at room temperature for 5 days followed by dissection.

### Preparation of SCRaMbLEd pool

The SCRaMbLE experiment using pCLB2-Cre-EBD-CYC1t was performed as described before^24^. Briefly, the yeast strains were transformed with the plasmid pCLB2-Cre-EBD-CYC1t, and selected on SC–His agar plates. The SCRaMbLE system was triggered upon β-estradiol. Yeast cells were then harvested by centrifugation at 2,000 ×g, washed twice with ddH_2_O to remove β-estradiol, resuspended in 50 mL of YPD liquid medium, and incubated for 24 h to generate the pool of SCRaMbLEd cells for further analysis.

### High-depth whole-genome sequencing

The next generation sequencing library was prepared using VAHTS Universal DNA Library Prep Kit for Illumina (Cat# ND607), following the manufacturer’s protocol with 1,000 ng input DNA. The prepared libraries were quantified by Qubit3.0 Fluorometer (Invitrogen, Carlsbad, CA, USA). These libraries with different indexes were multiplexed and loaded on an Illumina NovaSeq instrument according to manufacturer’s instructions (Illumina, San Diego, CA, USA). Sequencing was carried out using a 150 bp paired-end (PE) configuration; image analysis and base calling were conducted by the NovaSeq Control Software (HCS) + RTA 2.7 (Illumina) on the NovaSeq instrument. The sequencing libraries were made without PCR amplification to avoid creating artifact junctions driven by hybridization of loxPsym sites. Furthermore, in order to accurately estimate copy number of target regions, amplification-free sequencing was applied to decrease the likelihood that an appreciable proportion of these sequences would be duplicated and preserve a more even distribution of read coverage across the targeted sequencing regions.

### Analyses of novel junctions

Structural variations on the synthetic chromosomes were identified by the alignment of loxPsym sites and neighboring sequences to the yZSJ025 reference genome sequence. Reads containing loxPsym sites and their extensions by 116 bp on both sides were extracted. The following criteria were used to identify and screen rearrangements for further studies: (1) Reads containing the entire 34 bp loxPsym site sequences with flanking sequences belonging to two loxPsym sites of the reference; (2) Reads with one end less than 4 bp apart from loxPsym were excluded; (3) Reads containing two or more mismatched bases were excluded.

Identical reads were considered as a result of a single rearrangement event. Only two or more reads endorsing a rearrangement event were included in the further analyses.

### ATAC-seq

A total of ∼1,000,000 yeast cells were washed with 1 mL RSB buffer (10mM Tris-HCl pH 7.4, 10 mM NaCl, 3 mM MgCl_2_) once and incubated for 60 min at 37 [using zymolyase to digest cell walls (0.015 g/mL, Solarbio). Then the cell was resuspended in 1 mL lysis buffer (10 mM Tris-HCl pH 7.4, 10 mM NaCl, 3 mM MgCl_2_, 0.5% NP40, 0.1% Digitonin, 0.1% Tween-20, 1xProtease inhibitor) for 10 min at 4°C to lyse the yeast cell membrane to obtain the nucleus. Immediately following the nuclei prep, the pellet was resuspended in the Tn5 transposase reaction mix. The transposition reaction was carried out for 30 min at 37 °C. Tn5 transposed DNA were purified by AMPure DNA magnetic beads. A qPCR reaction was performed on a subset of the DNA to determine the optimum number of PCR cycles. The amplified libraries were run on an Agilent Tapestation 2100 (Agilent Technologies) to detect size distribution of library fragments. Biological replicates were performed in duplicate for all ATAC experiments. The final library was sequenced on an Illumina Nova-seq PE150 platform. Call peaks on replicates, self-pseudoreplicates using MACS2^54^ (--nomodel --extsize 200 --shift -100). The nucleoatac version 0.3.4 was used to call nucleosome positions and occupancy by ATAC data with default parameters.

### Analyses of the ATAC-seq signals

The average ATAC FPKM values of 90 hotspots, 90 coldspots and all 877 loxPsym sites in each bin were calculated. The mean ATAC-seq signals in the 400 bp surrounding hotspots and coldspots were normalized by the mean signal of all 877 loxPsym sites as follows:

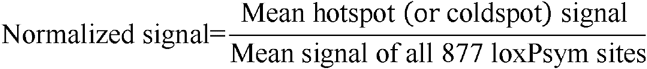

### Hi-C sequencing

The genomic DNA from exponential phase cells was cross-linked and digested with 200U *Mbo*I (NEB) as previously described^55^. Restriction fragment ends were labeled with biotinylated cytosine nucleotides by biotin-14-dCTP (TriLINK) followed by ligation. Purified DNA was sheared to a length of ∼400 bp. Point ligation junctions were pulled down with Dynabeads MyOne Streptavidin C1 (Thermofisher). The Hi-C library for Illumina sequencing was prepped by NEBNext Ultra II DNA library Prep Kit for Illumina (NEB Cat# E7645S) according to the manufacturers’ instructions. Fragments between 400 and 600 bp were paired-end sequenced on an Illumina Nova-seq PE150 platform.

### Construction of contact map and chromosome 3D model

After quality filtering using Trimmomatic (version 0.38), the clean Hi-C data two biological replicates for sample yZSJ025, was iteratively mapped to the yZSJ025 genome using the ICE software package (version 1f8815d0cc9e). Dandling ends and other unusable data were filtered, the valid pairs were used to analyze the correlation efficiency of the two biological replicates for each sample using QuASAR-Rep analysis (3DChromatin-ReplicateQC v0.0.1), then we pooled the data from two replicates together for further analysis. Valid pairs after pooling were binned into 1 Kb and 10 Kb nonoverlapping genomic intervals to generate contact maps. Raw Hi-C contact maps were normalized using iterative normalization method to eliminate systematic biases. The chromosomal 3D structure of the strain was inferred using the Pastis (v0.1) method^56^. with a multidimensional scaling (MDS) model. The 10-kb contact maps were used to construct the 3D model.

## Statistical analysis

Statistical analysis was performed using R (http://www.r-project.org/) or GraphPad Prism 8.0. Significance was determined via two-tailed, two-sample t tests. Pearson correlation analysis was applied to determine the correlation coefficient (r) and associated *p* values. A p value of ≤ 0.05 was considered statistically significant.

## Acknowledgments

We thank Jef D. Boeke from NYU Langone Health for many discussions and advice. We also thank Stephanie Lauer for helpful comments during manuscript preparation. We are grateful to Jef D. Boeke and Leslie A. Mitchell from NYU Langone Health, Huanming Yang, Yue Shen, Yun Wang and Tai Chen from BGI-Shenzhen, Yizhi Cai from the University of Manchester, for strain sharing and technical supports. This work is part of the Synthetic Yeast Genome Project (http://syntheticyeast.org/sc2-0/). This work was funded by the National Key Research and Development Program of China (2021YFC2100800) and the National Natural Science Foundation of China (21621004, 31861143017, 31971351). Fig.5 was created with BioRender.com.

## Author contributions

Conceptualization: YJY, YW; experiments: SZ, YZ, ZZ; analysis: YJY, YW, SZ, ZZ, LJ, LL; writing: YJY, YW, SZ, YZ, JT.

## Data and materials availability

Genome sequencing data have been submitted to NCBI Sequence Read Archive (SRA) under accession number PRJNA705059. The ATAC-seq and Hi-C sequencing data have been submitted to NCBI Gene Expression Omnibus (GEO) under accession number GSE168182. The ATAC-seq data of wild type *S. cerevisiae* is from GEO under accession number GSE66386.

## References

1. Good BH et al. The dynamics of molecular evolution over 60,000 generations. Nature 551, 45–50 (2017).

2. Jasinska W et al. Chromosomal barcoding of E. coli populations reveals lineage diversity dynamics at high resolution. Nat. Ecol. Evol. 4, 437–452 (2020).

3. Johnson MS, Martsul A, Kryazhimskiy S, Desai MM. Higher-fitness yeast genotypes are less robust to deleterious mutations. Science 366, 490–493 (2019).

4. Nguyen Ba AN et al. High-resolution lineage tracking reveals travelling wave of adaptation in laboratory yeast. Nature 575, 494–499 (2019).

5. Peter J, Schacherer J. Population genomics of yeasts: towards a comprehensive view across a broad evolutionary scale. Yeast 33, 73–81 (2016).

6. Gallone B et al. Domestication and divergence of Saccharomyces cerevisiae beer yeasts. Cell 166, 1397–1410 (2016).

7. Botstein D, Fink GR. Yeast: an experimental organism for 21st century biology. Genetics 189, 695–704 (2011).

8. Peter J et al. Genome evolution across 1,011 Saccharomyces cerevisiae isolates. Nature 556, 339–344 (2018).

9. Shen X et al. Tempo and mode of genome evolution in the budding yeast subphylum. Cell 175, 1533–1545 (2018).

10. Marsit S et al. Evolutionary biology through the lens of budding yeast comparative genomics. Nat. Rev. Genet. 18, 581–598 (2017).

11. Yue J et al. Contrasting evolutionary genome dynamics between domesticated and wild yeasts. Nat. Genet. 49, 913–924 (2017).

12. Dujon B. Yeast evolutionary genomics. Nat. Rev. Genet. 11, 512–524 (2010).

13. Hutchison CA et al. Design and synthesis of a minimal bacterial genome. Science 351, d6253 (2016).

14. Ostrov N et al. Design, synthesis, and testing toward a 57-codon genome. Science 353, 819–822 (2016).

15. Fredens J et al. Total synthesis of Escherichia coli with a recoded genome. Nature 569, 514–518 (2019).

16. Richardson SM et al. Design of a synthetic yeast genome. Science 355, 1040–1044 (2017).

17. Chen W et al. An artificial chromosome for data storage. Natl Sci Rev 8, b28 (2021).

18. Lu X, Ellis T. Self-replicating digital data storage with synthetic chromosomes. Natl Sci Rev 8 (2021).

19. Dymond JS et al. Synthetic chromosome arms function in yeast and generate phenotypic diversity by design. Nature 477, 471–476 (2011).

20. Blount BA et al. Rapid host strain improvement by in vivo rearrangement of a synthetic yeast chromosome. Nat Commun 9, 1932 (2018).

21. Jia B et al. Precise control of SCRaMbLE in synthetic haploid and diploid yeast. Nat Commun 9, 1–13 (2018).

22. Liu W et al. Rapid pathway prototyping and engineering using in vitro and in vivo synthetic genome SCRaMbLE-in methods. Nat Commun 9, 1936 (2018).

23. Luo Z et al. Identifying and characterizing SCRaMbLEd synthetic yeast using ReSCuES. Nat Commun 9, 1–10 (2018).

24. Shen MJ et al. Heterozygous diploid and interspecies SCRaMbLEing. Nat Commun 9, 1–8 (2018).

25. Shen Y et al. SCRaMbLE generates designed combinatorial stochastic diversity in synthetic chromosomes. Genome Res. 26, 36–49 (2016).

26. Wang J et al. Ring synthetic chromosome V SCRaMbLE. Nat Commun 9, 3783 (2018).

27. Wu Y et al. In vitro DNA SCRaMbLE. Nat Commun 9, 1935–1939 (2018).

28. Mitchell LA et al. Synthesis, debugging, and effects of synthetic chromosome consolidation: synVI and beyond. Science 355, f4831 (2017).

29. Shen Y et al. Deep functional analysis of synII, a 770-kilobase synthetic yeast chromosome. Science 355, f4791 (2017).

30. Wu Y et al. Bug mapping and fitness testing of chemically synthesized chromosome X. Science 355, f4706 (2017).

31. Xie Z et al. “Perfect” designer chromosome V and behavior of a ring derivative. Science 355, f4704 (2017).

32. Annaluru N et al. Total synthesis of a functional designer eukaryotic chromosome. Science 344, 55–58 (2014).

33. Lin Q, Qi H, Wu Y, Yuan Y. Robust orthogonal recombination system for versatile genomic elements rearrangement in yeast Saccharomyces cerevisiae. Sci Rep-Uk 5, 15249 (2015).

34. Karimova M et al. Vika/vox, a novel efficient and specific Cre/loxP-like site-specific recombination system. Nucleic Acids Res. 41, e37 (2013).

35. Schep AN et al. Structured nucleosome fingerprints enable high-resolution mapping of chromatin architecture within regulatory regions. Genome Res. 25, 1757–1770 (2015).

36. Duan Z et al. A three-dimensional model of the yeast genome. Nature 465, 363–367 (2010).

37. Mercy G et al. 3D organization of synthetic and scrambled chromosomes. Science 355, f4597 (2017).

38. Wang P et al. SCRaMbLEing of a synthetic yeast chromosome with clustered essential genes reveals synthetic lethal interactions. Acs Synth Biol 9, 1181–1189 (2020).

39. Guo F, Gopaul DN, Van Duyne GD. Structure of Cre recombinase complexed with DNA in a site-specific recombination synapse. Nature 389, 40–46 (1997).

40. Ames RM et al. Gene duplication and environmental adaptation within yeast populations. Genome Biol Evol 2, 591–601 (2010).

41. Bergström A et al. A high-definition view of functional genetic variation from natural yeast genomes. Mol. Biol. Evol. 31, 872–888 (2014).

42. Brown CA, Murray AW, Verstrepen KJ. Rapid expansion and functional divergence of subtelomeric gene families in yeasts. Curr. Biol. 20, 895–903 (2010).

43. Parker R. RNA degradation in Saccharomyces cerevisiae. Genetics 191, 671–702 (2012).

44. Yadav V, Sun S, Coelho MA, Heitman J. Centromere scission drives chromosome shuffling and reproductive isolation. P. Natl Acad. Sci. Usa 117, 7917–7928 (2020).

45. Guin K et al. Spatial inter-centromeric interactions facilitated the emergence of evolutionary new centromeres. Elife 9, e58556 (2020).

46. Sankaranarayanan SR et al. Loss of centromere function drives karyotype evolution in closely related Malassezia species. Elife 9, e53944 (2020).

47. Hadi K et al. Distinct Classes of Complex Structural Variation Uncovered across Thousands of Cancer Genome Graphs. Cell 183, 197–210 (2020).

48. Zhang Y et al. Spatial organization of the mouse genome and its role in recurrent chromosomal translocations. Cell 148, 908–921 (2012).

49. Fudenberg G, Getz G, Meyerson M, Mirny LA. High order chromatin architecture shapes the landscape of chromosomal alterations in cancer. Nat. Biotechnol. 29, 1109–1113 (2011).

50. Pinglay S et al. Synthetic genomic reconstitution reveals principles of mammalian Hox cluster regulation. bioRxiv, 2021-2027 (2021).

51. Zhao Y et al. Debugging and consolidating multiple synthetic chromosomes reveals combinatorial genetic interactions. bioRxiv, 2022-2024 (2022).

52. Gibson DG et al. Enzymatic assembly of DNA molecules up to several hundred kilobases. Nat. Methods 6, 343–345 (2009).

53. Gietz RD, Schiestl RHW, A. R. Woods R. Studies on the transformation of intact yeast cells by the LiAc/SS-DNA/PEG procedure. Yeast 11, 355–360 (1995).

54. Zhang Y et al. Model-based Analysis of ChIP-Seq (MACS). Genome Biol. 9, R137 (2008).

55. Lieberman-Aiden E et al. Comprehensive mapping of long-gange interactions reveals folding principles of the human genome. Science 326, 289–293 (2009).

56. Varoquaux N, Ay F, Noble WS, Vert JP. A statistical approach for inferring the 3D structure of the genome. Bioinformatics 30, i26–i33 (2014).

