## Supplemental figure for "Synthetic genome rearrangement reveals dynamics of chromosome evolution shaped by hierarchical chromatin organization" (from another version of this preprint: version 3, 2022-06-29)

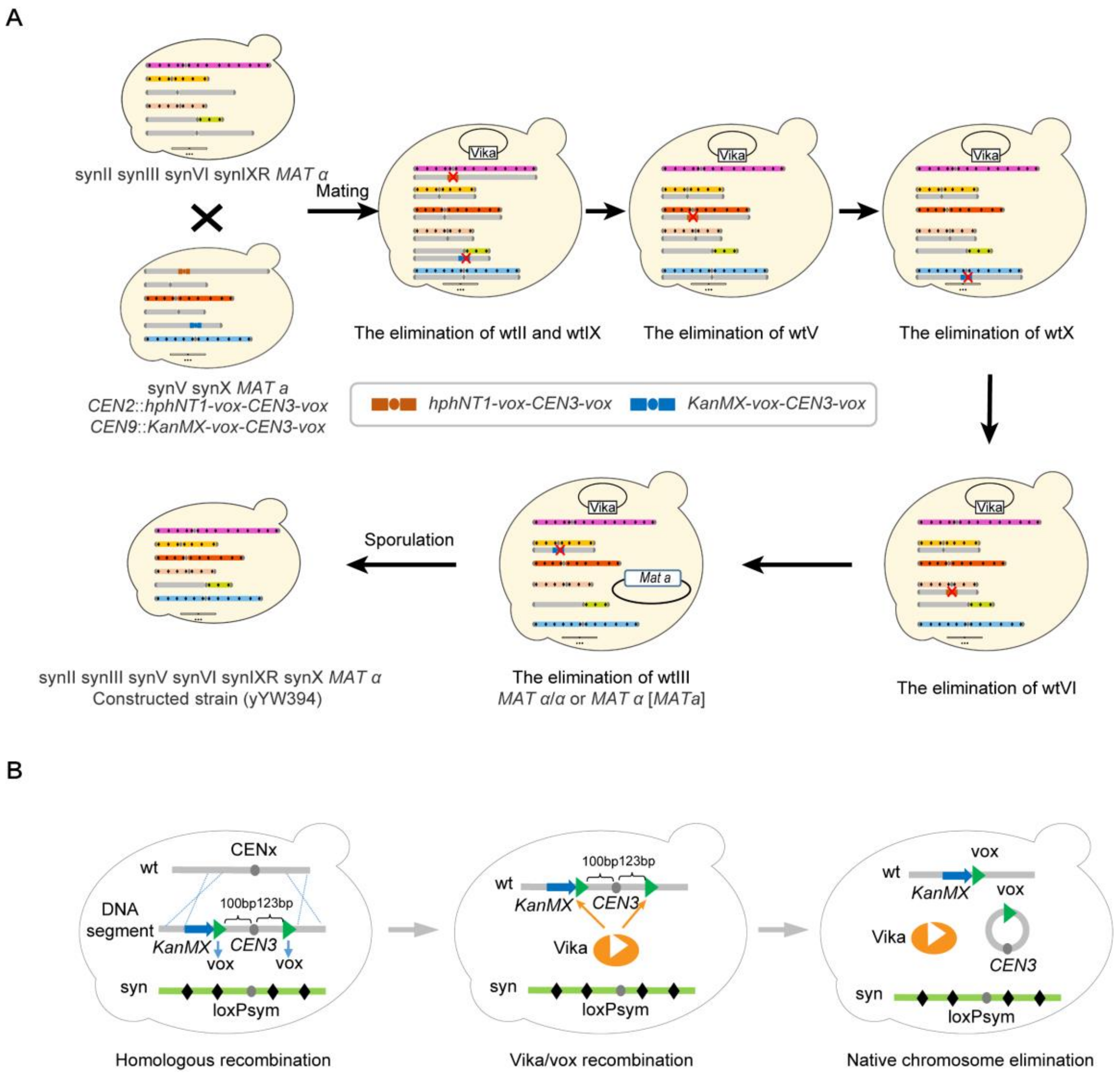


**Figure S1.** Consolidation and adaptive laboratory evolution of a poly-synthetic chromosome strain. (**A**) *KanMX-vox-CEN3-vox* and *hphNT1-vox-CEN3-vox* constructs were integrated into the centromere regions of native chromosomes II and IX in strain synV, synX, which were then mating with strain synII, III, VI, IXR. When Vika recombinase was induced, the native chromosomes were eliminated and confirmed by PCRTag analysis. Following the same strategy, wtV, wtX, wtVI, and wtIII were eliminated sequentially. An episomal copy of *MATa* was introduced to permit sporulation after elimination. Haploid synthetic chromosome strain (yYW394) was then generated by sporulation. Synthetic chromosomes were labeled with different colors and the native chromosomes were shown in grey. (**B**) Mechanism for chromosome elimination Vika/vox. The centromere constructs containing two vox, *CEN3* and *KanMX* or *hphNT1*. The constructs were integrated into the centromere regions of native chromosomes.

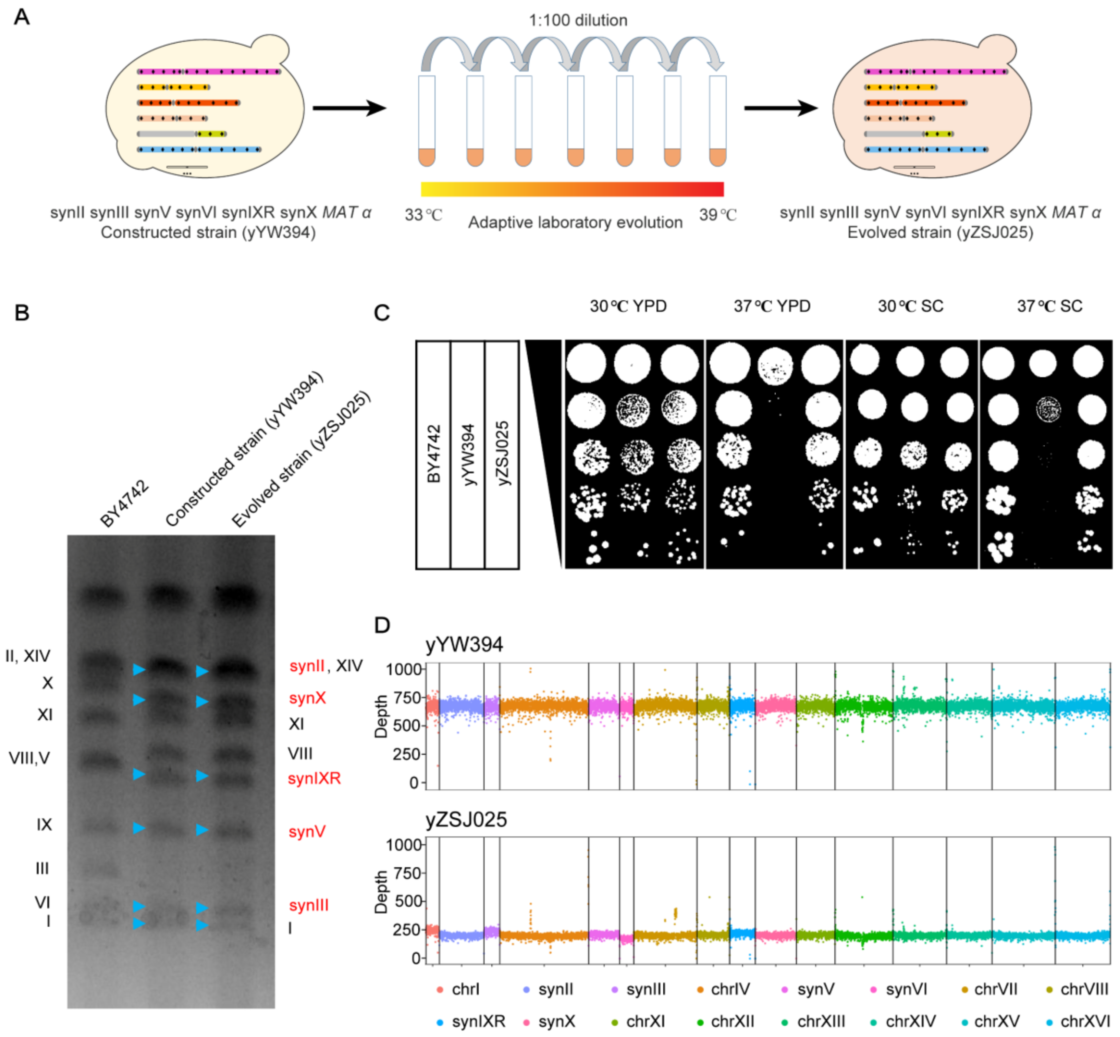


**Figure S2.** The phenotype and genotype of yYW394 and yZSJ025. (**A**) Adaptive laboratory evolution of yYW394 was carried out from 33°C to 39°C to generate an evolutionary haploid strain (yZSJ025). Synthetic chromosomes were labeled with different colors and the native chromosomes were shown in grey. (**B**) Karyotypic analysis of BY4742, yYW394 and yZSJ025 by pulsed-field gel electrophoresis; note that IXL-synIXR and native IX migrate identically. (**C**) Ten-fold serial dilutions of BY4742, yYW394 and yZSJ025 on various types of media at 30°C or 37°C. YPD, yeast extract peptone dextrose; SC, synthetic complete medium. (**D**) The coverage maps of yYW394 and yZSJ025. The sequencing depths of all sixteen chromosomes were at the same scale. No major structural variations were detected in either strain. Only 25 minor sequence variations including SNP and short indel (Table S2) occurred in yYW394 compared to previously reported individual synthetic chromosomes, and 37 additional mutations developed in yZSJ025 (Table S3). SynVI was tentatively identified as a ring chromosome.

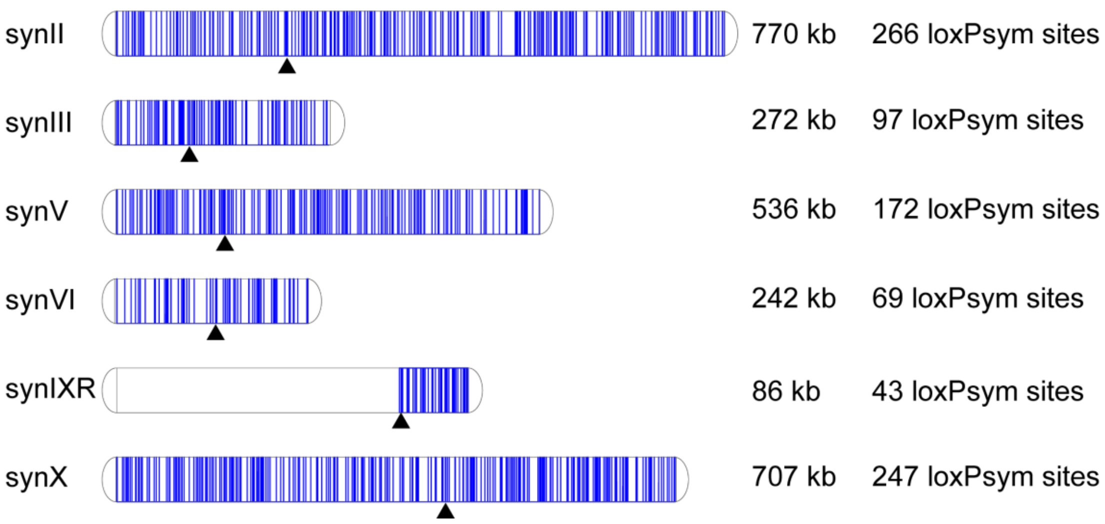


**Figure S3.** The distributions of 894 loxPsym sites.

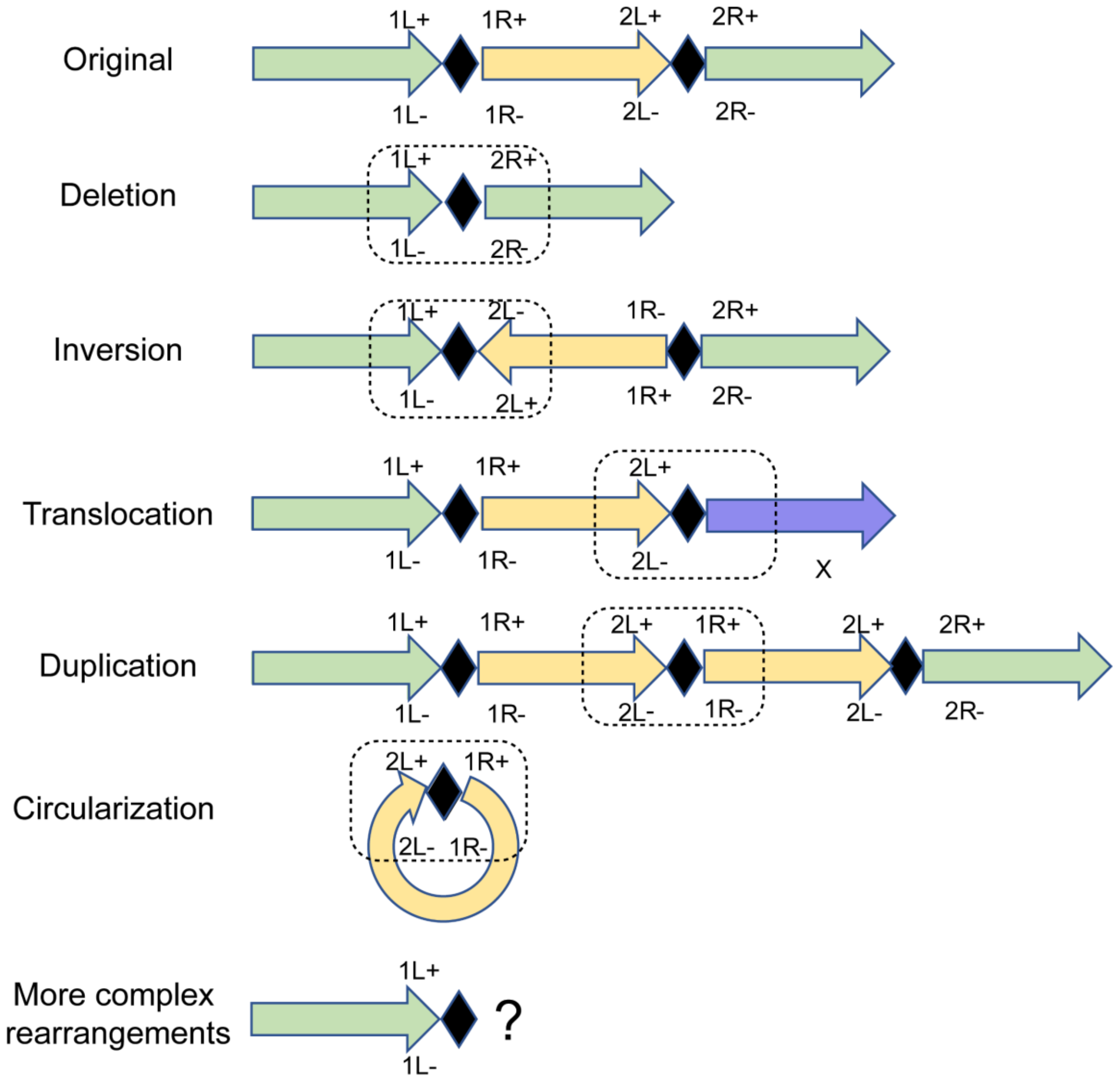


**Figure S4.** Classification strategy for novel junction analysis. “1” and “2” indicates the order of segment in a synthetic chromosome; “X” indicates the segment in another synthetic chromosome; “+” and “-” indicate the positive and negative strands. Deletion events could be identified as a “1L+ & 2R+” or “1L- & 2R-” connection type, inversion events could be identified as a “1L+ & 2L-” or “1L- & 2L+” connection type, and translocation events could be identified as a “2L+ & X” or “2L- & X” connection type. Other types of junctions could be identified as evidence of complex events, such as (duplication and circularization: “2L+ & 1R+” or “2L- & 1R-”).

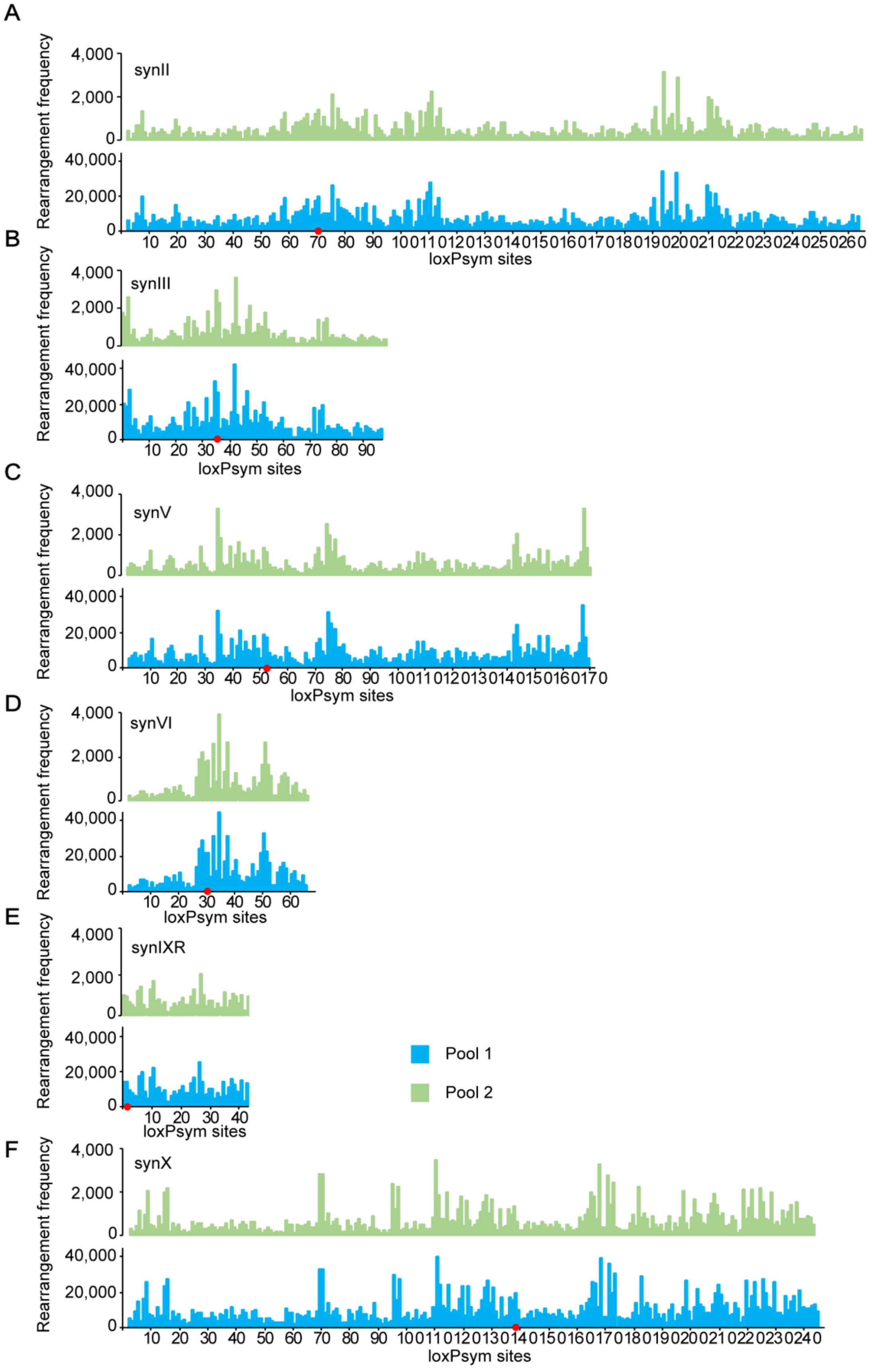


**Figure S5.** Landscape of the rearrangement frequencies along synII, synIII, synV, synVI, synIXR, and synX in biological replicate. Red dots indicate centromeres.

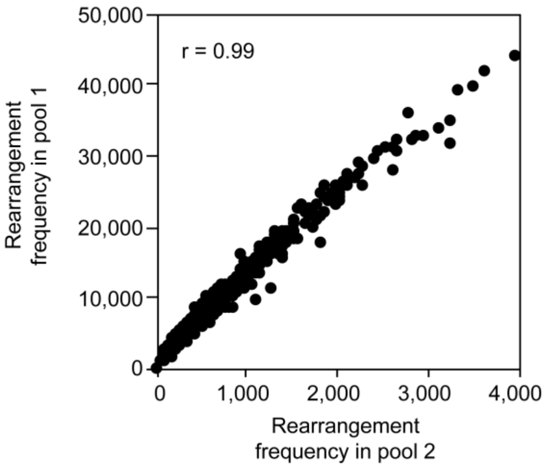


**Figure S6.** Comparison of rearrangement frequencies in the replicates of SCRaMbLEd cells (r = 0.99, $p = 1.7\times10^{-232}$). Pearson correlation analysis was applied to determine the correlation coefficient and associated p values.

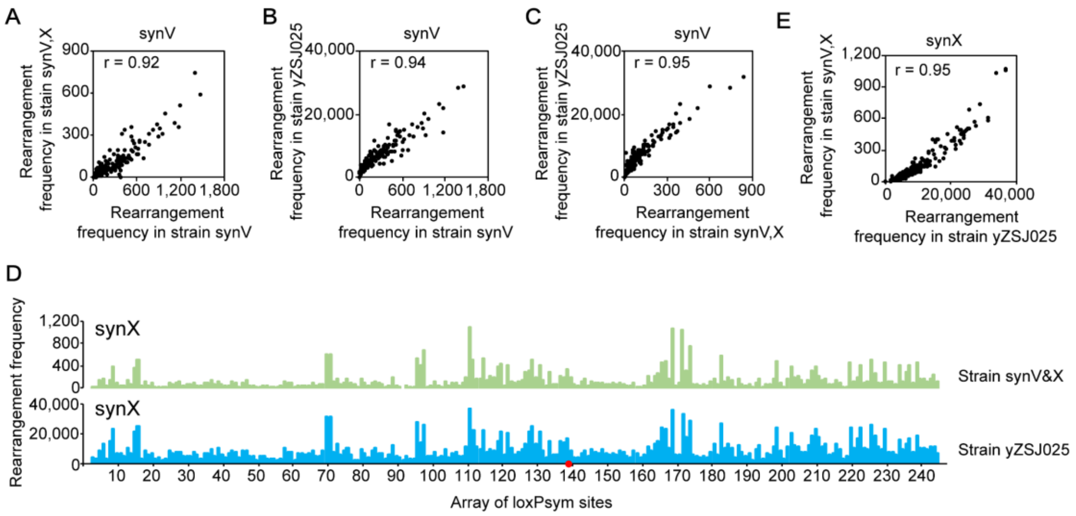


**Figure S7.** Intra-chromosomal rearrangement patterns of synV and synX. (**A-C**) Comparisons of rearrangement frequencies in synV following SCRaMbLE of strains yXZX846 (synV), yYW169 (synV, synX) and yZSJ025. (**A**) Comparisons of strains yXZX846 and yYW169 (r = 0.92, $p = 5.1\times10^{-12}$). (**B**) Comparisons of strains yXZX846 and yZSJ025 (r = 0.94, $p = 6.0\times10^{-49}$). (**C**) Comparisons of strains yYW169 and yZSJ025 (r = 0.95, $p = 3.9\times10^{-51}$). (**D**) Intra-chromosomal rearrangement patterns of synX following SCRaMbLE in two different yeast strains, yYW169 and yZSJ025. The red dots indicate centromeres. (**E**) Comparisons of rearrangement frequencies in synX following SCRaMbLE of strains yYW169 and yZSJ025 (r = 0.95, $p = 1.6\times10^{-71}$). Pearson correlation analysis was applied to determine the correlation coefficient and associated p values.

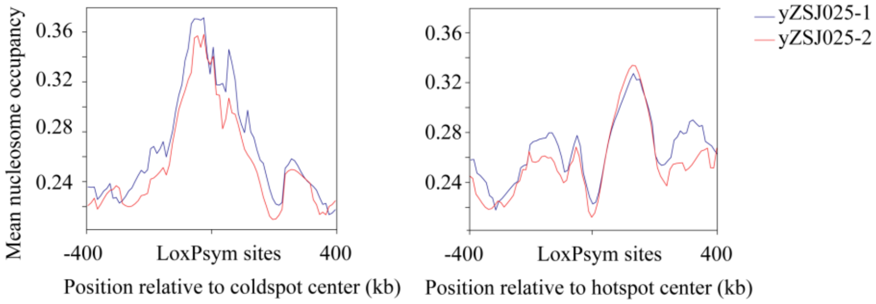


**Figure S8.** Mean nucleosome occupancy around hotspots and coldspots.

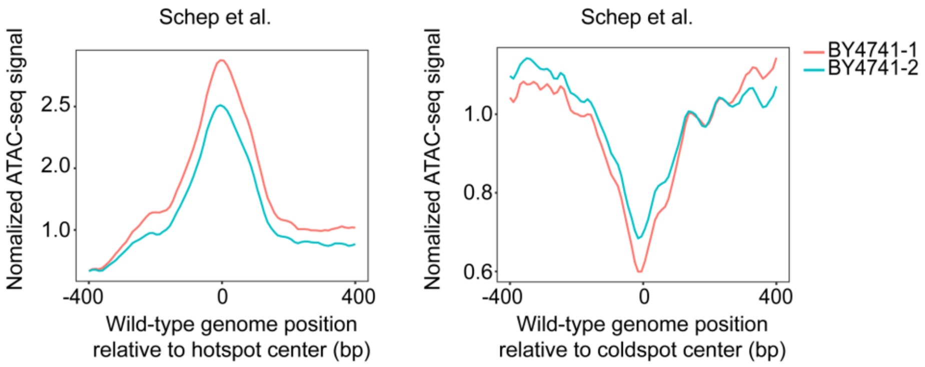


**Figure S9.** Normalized mean ATAC-seq signals around the wide-type matched positions of hotspots and coldspots.

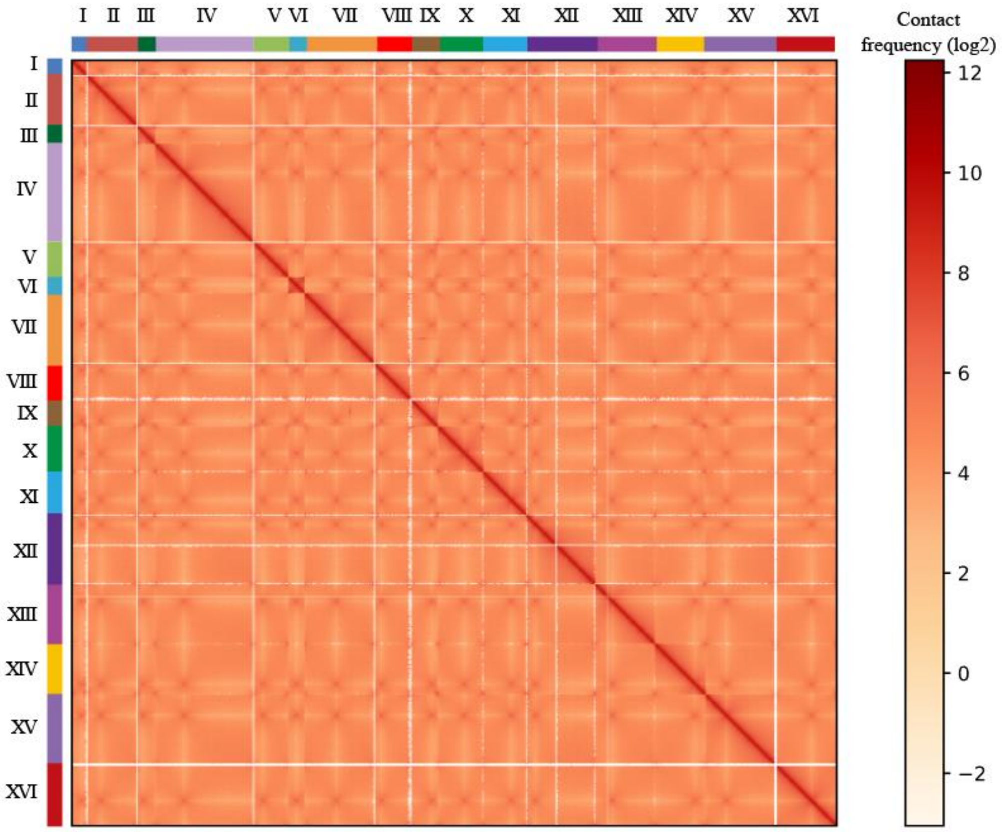


**Figure S10.** Contact map of yZSJ025. A normalized contact map of all chromosomes of yZSJ025 is shown (bin size, 10 kb). The normalized contact frequencies are indicated on a log2 scale with colors ranging from light yellow (few contacts) to red (many contacts).

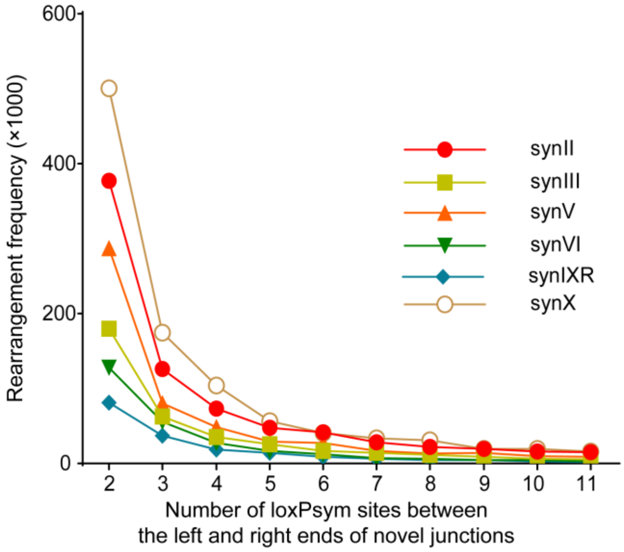


**Figure S11.** Distributions of the numbers of loxPsym sites between the left and right ends of novel junctions for all synthetic chromosomes.
